# Multi◻region whole◻genome and transcriptomic profiling uncovers plastic, subclone◻linked cell states in high◻grade diffuse astrocytomas

**DOI:** 10.64898/2026.08.11.743949

**Authors:** Serafiina Ohlsbom, Sonja Mäntylä, Reetta Nätkin, Ismaïl Hermelo, Anssi Nurminen, Aliisa M. Tiihonen, Iida Salonen, Elisa Vuorinen, Kristiina Nordfors, Hannu Haapasalo, Kirsi J. Rautajoki, Joonas Haapasalo, Matti Nykter

## Abstract

Intratumoral heterogeneity is a defining feature of high-grade astrocytomas and a major contributor to treatment resistance. Yet how genomic diversification intersects with transcriptional plasticity remains incompletely understood. We performed high-resolution multi-omic profiling of three complex, treatment-naïve tumors (two IDH-wildtype glioblastomas and one IDH-mutant grade 4 astrocytoma). By integrating whole-genome sequencing (WGS), bulk and single-cell RNA sequencing (scRNA-seq), and histopathology across four anatomically distinct regions per tumor, we mapped the co-evolution of genome and transcriptome. Despite striking regional differences in morphology and cellular states, genomic evolution was predominantly trunk-dominated. Most driver alterations were clonal across regions, indicating early acquisition and stable genomic backbones. The IDH-mutant tumor showed linear evolution with localized hypermutation, whereas glioblastomas displayed modest late-branching subclones. In contrast, transcriptional heterogeneity was pronounced and spatially structured. Distinct genetic subclones preferentially occupied divergent transcriptional states. However, subclones shared across regions frequently adopted different phenotypes depending on local microenvironment. Single-cell reconstruction from matched patient-derived cell lines resolved subclone-associated trajectories, revealing dynamic transitions between proliferative and inflammatory states. This study provides a framework for understanding how early-established genomic backbones and regional transcriptional plasticity jointly drive phenotypic diversity. While single biopsies may capture truncal drivers, resolving clinically relevant heterogeneity requires multi-region and single-cell approaches.

## Introduction

Adult-type high-grade astrocytomas, especially glioblastomas, are characterized by pronounced intratumoral heterogeneity. It is associated with decreased survival^1^ and suggested as a cause of treatment failure and therapy resistance^2,3^. The striking morphologic and clinical diversity arises from layered biological heterogeneity encompassing genetic, transcriptional, epigenetic, histological, and microenvironmental differences^2,4^, collectively modulating tumor behavior, therapeutic response, and progression. Genomic heterogeneity reflects ongoing accumulation of stable, usually irreversible, alterations and chromosomal instability shaped by clonal evolution, whereas transcriptomic and epigenetic heterogeneity captures more dynamic processes, including differential expression, epigenomic remodeling, chromatin organization, and cancer cell plasticity^2^. Tumor formation is further influenced by microenvironmental heterogeneity, in which immune, stromal, and vascular cells co-evolve with tumor cells under local selection pressures^2,4^.

Traditional genomic diffuse glioma studies have focused on characterizing and classifying tumors based on common molecular alterations observed^5–7^, relying on a single representative biopsy per tumor. While such approaches capture driver events, they fail to resolve how tumors evolve in anatomical context. Multi-regional genomic studies have addressed these caveats but remain limited in the breadth and depth needed to map full subclonal architecture, for example due to the lack of whole-genome coverage^8–14^. Single-cell and spatial transcriptomic studies have revealed that gliomas contain multiple cancer cell states^1,15,16^ with the capacity to transit between phenotypes^15,17^, but these current studies are limited in the ability to link transcriptional states to evolutionary lineages.

Although key genetic alterations in high-grade astrocytomas are well characterized, the spatial relationships among genetic diversification, transcriptional plasticity, and microenvironmental context remain incompletely understood. Moreover, the extent to which subclonal diversity contributes to regional phenotypes is unresolved. Understanding how specific mutations map onto transcriptomic and microenvironmental programs is critical for defining how gliomas grow, invade, and escape therapy. To establish a framework for the dissection and integration across these interconnected layers of heterogeneity at a genome-wide scale, we performed a pilot study with deep multi-omic profiling of three spatially complex high-grade diffuse astrocytoma tumors—two glioblastomas and one IDH-mutant grade 4 astrocytoma. To interrogate how genetic and plastic heterogeneity interact across tumor anatomy, and to evaluate the implications of these processes for clinical actionability and biopsy fidelity, we integrated whole-genome sequencing, bulk RNA-seq, scRNA-seq of patient-derived cell lines, and histopathology across four spatially distinct regions per tumor.

## Results

### Spatially sampled tumor regions exhibit marked histopathological diversity and a shared genomic backbone

We selected three patients with large, aggressive, high-grade primary diffuse astrocytoma tumors based on their size and morphological complexity to enable detailed investigation of intratumoral heterogeneity and tumor evolution (Supplementary Fig. 1). One tumor was diagnosed as IDH-mutant grade 4 astrocytoma (DA1, age-group 25–35 years), and two as IDH-wildtype glioblastomas (GB1, 55–65 years, and GB2, 65–75 years) (Supplementary Table 1). We collected samples from the tumor-normal interface (R1), progressing to deeper intratumoral sites (R2 and R3), and toward the opposite tumor-normal interface (R4), allowing for comparisons between peritumoral region and tumor core (Fig. 1a). All tumors were located in the left hemisphere, in temporal (DA1) or frontal (GB1 and GB2) lobes (Fig. 1b). Clinical outcomes diverged markedly: DA1 remains relapse-free six years after subtotal resection and chemoradiotherapy, exceeding typical outcomes^18^, whereas GB1 and GB2 showed much poorer outcomes of 10 and 3 months, respectively, which are shorter than the typical 12–15 months reported for glioblastoma^19,20^.

**Fig. 1.**
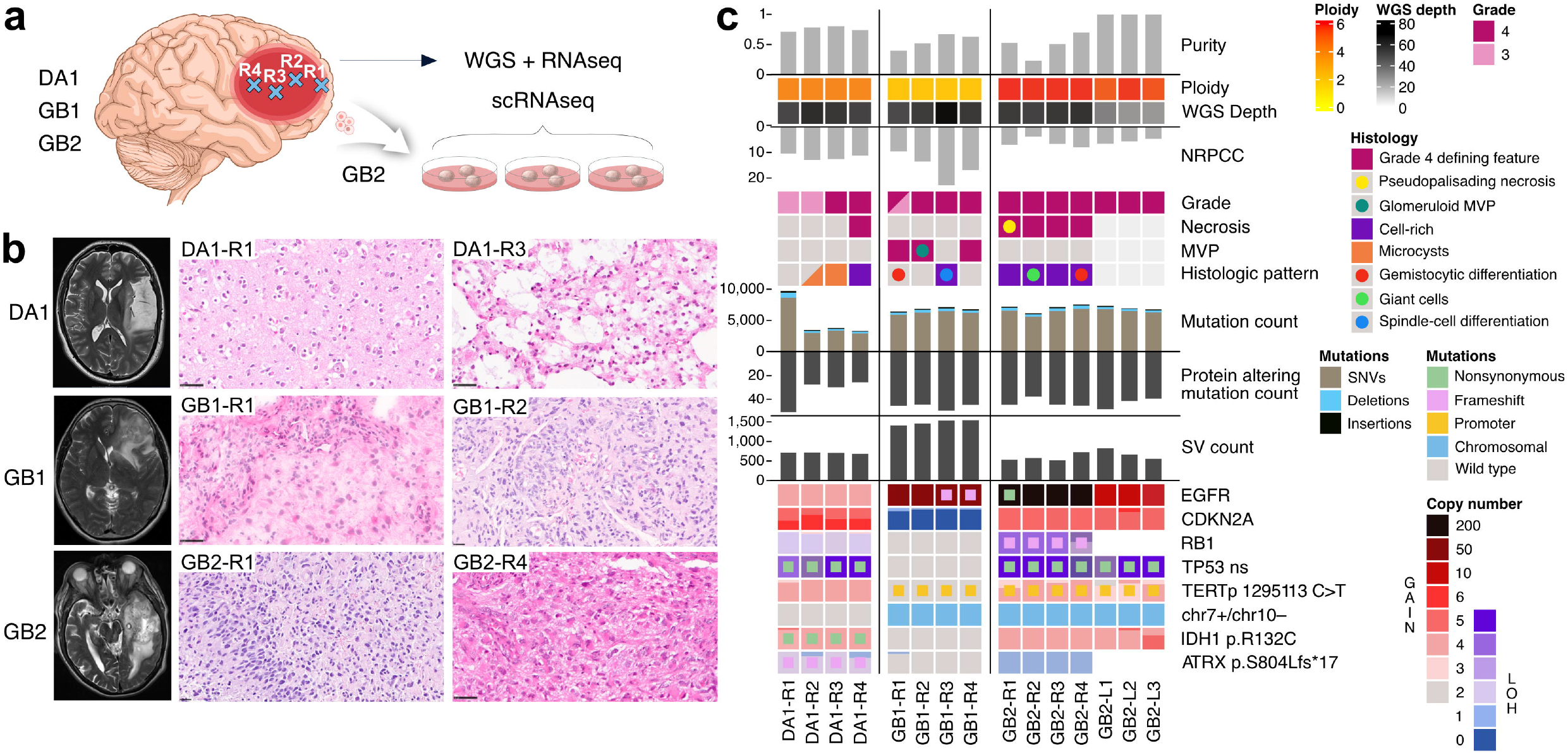
Multi-regional sampling reveals histopathological diversity and distinct genomic architectures across astrocytoma and glioblastoma tumors. **a)** Three large diffuse astrocytomas (two glioblastomas and one IDH-mutant astrocytoma; GB1, GB2, and DA1) were selected for assessing intratumoral heterogeneity and tumor evolution by sampling four spatially distinct regions from each tumor (illustrative). In addition, three patient-derived cell lines were created from two GB2 tumor regions. **b)** H&E images highlight oligodendroglioma-like differentiation and microcysts in grade 3 (R1) and grade 4 (R3) DA1 regions, respectively. Tumor border of high grade and lower grade differentiation (R1, fresh frozen tissue) and glomeruloid MVP (R2) were detected in GB1, and pseudopalisading necrosis (R1) and gemistocytic differentiation (R4) in GB2 regions. Magnification 40x. **c)** The tumors exhibited the characteristic molecular alterations of glioblastoma and IDH-mutant astrocytoma but showed distinct genomic profiles: DA1 displayed a tetraploid genome and regional hypermutation; GB1 had a diploid genome with a high number of SVs; and GB2 carried a TP53 mutation, showed extensive aneuploidy, and harbored extreme EGFR amplification. For subclonal CNAs, the clone proportions are shown in square height proportions. ns: nonsynonymous, fs: frameshift.

Histological heterogeneity was striking both within and between tumors. DA1 regions R1 and R2 represented typical grade 3 morphology with oligodendroglioma-like differentiation and small microcystic features (R2), while R3 and R4 showed grade 4 morphology with highly prevalent microcysts (R3) and focal necrosis (R4) (Fig. 1b–c, Supplementary Fig. 2a). Driver mutations in *IDH1* (p.R132C), *ATRX* (p.S804Lfs*17), and *TP53* (p.K132E) across all regions confirmed the molecular diagnosis (Supplementary Table 2, Supplementary Fig. 3). In contrast, both glioblastomas GB1 and GB2 were histologically uniformly grade 4 but showed phenotypically diverse morphologies. GB1-R1 captured the tumor border of low- and high-grade differentiation together with clear microvascular proliferation (MVP) and gemistocytic-like astrocytoma cells, while deeper regions represented more cell-rich morphology with glomeruloid MVP (GB1-R2) and spindle cell differentiation (GB1-R3). GB2 appeared histologically more aggressive than GB1, with cell-rich morphology and wide necrosis throughout the tumor. GB2 margin (GB2-R1) exhibited a tumor core feature pseudopalisading necrosis^21^, while deeper regions showed several giant cells (GB2-R2) and gemistocytic differentiation (GB2-R4). Both glioblastomas carried *EGFR* amplification and *TERT* promoter mutations in all regions, while *CDKN2A* deletion was only detected in GB1 and *TP53* (p.R248Q) mutation in GB2 (Supplementary Tables 3–4). The proliferative index measured by Ki-67 staining was low in DA1 (10%), moderate in GB1 (25%), and high in GB2 (40%) (Supplementary Fig. 2b).

In DA1 samples, WGS revealed tetraploid tumor genomes with minimal regional variation (3.72–3.76) and high tumor purity (70–88%), reflecting a tumor-cell-dominant tissue composition (Fig. 1c, Supplementary Fig. 4) (Supplementary Table 1). In contrast, GB1 and GB2 samples showed lower and more variable purity estimates (40–67% in GB1 and 23–67% in GB2, Supplementary Fig. 5–6), highlighting the complex, infiltrative tissue architecture underlying glioblastoma heterogeneity. GB1 tumor genomes were diploid. Conversely, GB2 tumor cells displayed evidence of high polyploidy, which was confirmed by karyotype analysis of three GB2 patient-derived cell lines, revealing an average of 116 chromosomes per cell in 15 analyzed cells (Supplementary Table 5), consistent with an approximate ploidy of five. This was comparable with WGS estimates of 4.66–5.69. Due to high ploidy in GB2, number of reads per chromosome copies (NRPCC) remained low (<10) in all GB2 regions (Fig. 1c), resulting in noisier subclonal variant detection than in DA1 or GB1.

### Extensive evolutionary trunks with late-arising spatial genomic divergence

Understanding whether the marked clinical and histological heterogeneity of diffuse astrocytomas arises from genomic diversification or later, plastic processes requires resolving how evolution unfolds across tumor space. To assess this, we leveraged the multi-region WGS to profile somatic alterations and to reconstruct subclonal structure at peritumoral and intratumoral sites.

The IDH-mutant DA1 exhibited early loss of heterozygosity (LOH) on chromosomes 4q, 11p, 13q, and 17p (affecting e.g. *RB1*, *BRCA2*, and *TP53*), followed by *TP53* locus duplication, whole-genome duplication (WGD) and a chromothripsis involving chromosomes 2 and X (Fig. 2a, Supplementary Fig. 4 and 7, Supplementary Tables 6–8). These early events were clonal across all regions (R1-R4). Truncal *ATRX* frameshift (p.S804Lfs*17) (Supplementary Fig. 3) and *TP53* mutation (p.K132E) (Supplementary Table 2) showed high variant allele fractions (VAF) (0.79–0.85 for *ATRX*; 0.81–0.92 for TP53), indicating early acquisition prior to segmental duplication and/or WGD. Single nucleotide variant (SNV) clustering supported a predominantly linear tumor evolution with a clonal trunk comprising 1960 SNVs (22% of 9111 total SNVs) (Supplementary Fig. 8). In contrast to deeper tumor core regions, DA1-R1 showed notable accumulation of private SNVs (4806, 53%) and indels, consistent with localized hypermutation driven predominantly by clock-like SBS5 mutational processes (Supplementary Fig. 9). No direct driver for hypermutation phenotype was observed, but 268–501bp deletions upstream of *POLE* (777bp and 5656bp) were detected in all regions (Supplementary Table 8), however, the RNA expression was not affected (Supplementary Fig. 10). Interestingly, region R1 showed grade 3 histology with only mild atypia (Fig. 1c), suggesting that the localized hypermutation reflects ongoing evolutionary activity that may precede detectable phenotypic divergence. Regions R1 and R3 shared a subclone (orange) containing 400 SNVs present in 40% and 37% of cancer cells (Fig. 2a, Supplementary Fig. 8), which harbored an *ATXN1*–*CA10* fusion (Supplementary Table 8); *CA10* encodes a carbonic anhydrase-related protein shown to suppress neuronal activity-dependent glioma growth^22^ and showed slightly reduced expression in affected regions compared to others. While a marked histological diversity was observed across regions R1–R4 (Fig. 1c, Supplementary Fig. 2), DA1 exhibited early shared genomic alterations followed by comparatively more recent, region-restricted hypermutation, indicating linear evolution with moderate spatial genomic divergence.

**Fig. 2.**
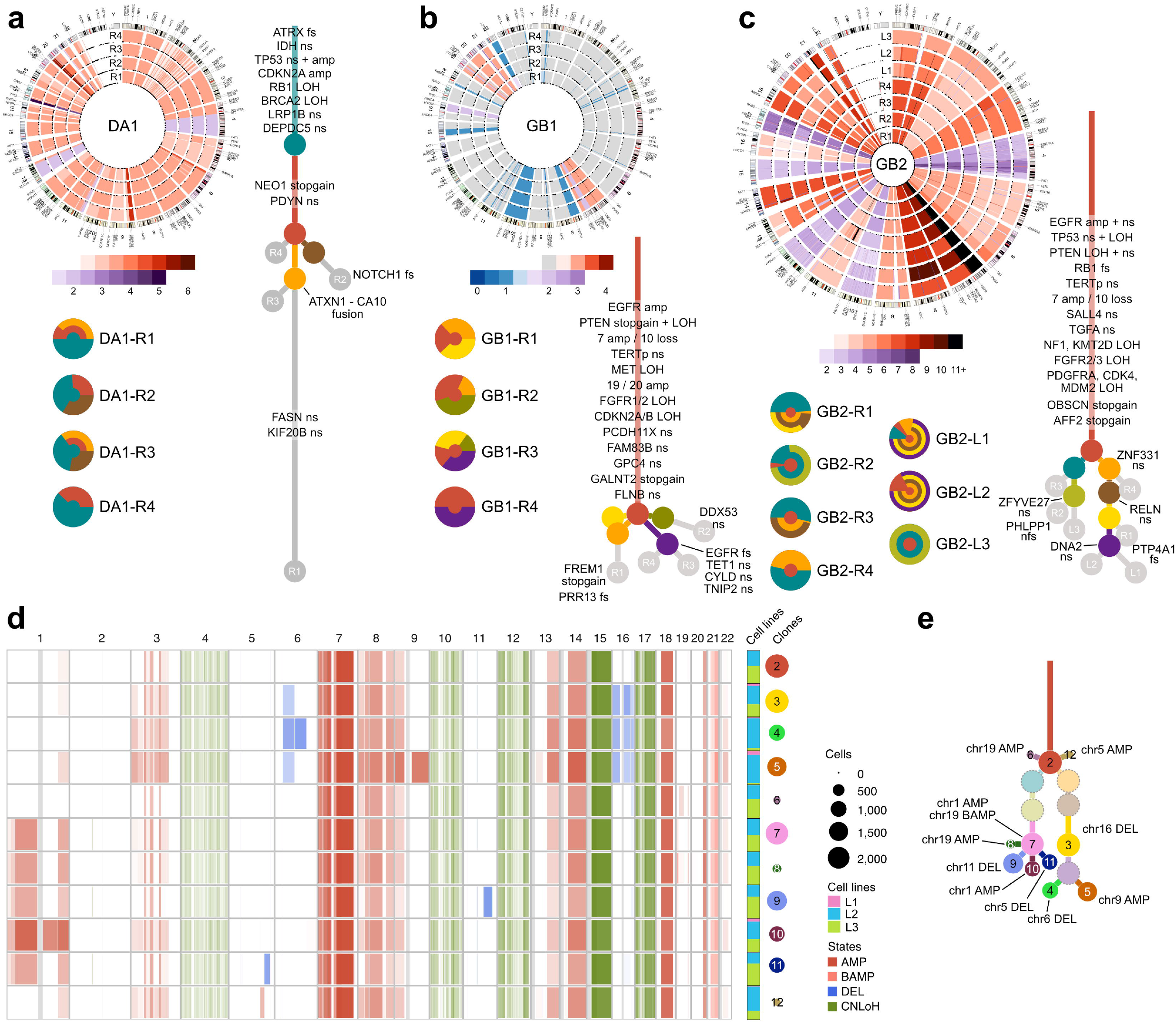
Copy number profiles and phylogenetic reconstruction define shared evolutionary trunks and modest spatial genomic divergence. **a)** CNAs across IDH-mutant DA1 tumor regions and chromosomes in circos plot show clonal WGD and chromothripsis event affecting e.g. DNA repair genes *MSH2* and *MSH6* loci in chr2. The evolutionary tree depicts a regional hypermutation in DA1-R1. Jawbreaker plots show the subclonal compositions of each tumor region. **b)** GB1 trunk included typical glioblastoma alterations such as *EGFR* amplification, full inactivation of *PTEN*, and *TERT* promoter mutation (C>T at chr5:1295113). The purple clone shared by GB1-R3–R4 contained e.g. *EGFR* frameshift (p.I1018Pfs*2), while the orange clone in regions R1–R2 harbored subclonal copy number losses. **c)** In addition to traditional glioblastoma oncogenes and tumor suppressors, GB2 harbored a truncal alteration in *TGFA* (p.R81C), an EGFR ligand. The purple subclone observed only in cell lines L1 and L2 contained a nonsynonymous mutation in DNA replication gene *DNA2* (p.P891S). For each tumor, evolutionary tree branch lengths are scaled by the number of SNVs. Private mutations in gray clones are placed in the branch with the highest cancer cell fraction. **d)** GB2-L1–L3 scRNA-seq copy number alterations show shared subclones between tumor regions and clones specific to L2 cell line. **e)** scRNA-seq copy number derived clones are separated into two main branches (clones 3 and 7), similar to the evolutionary tree based on GB2 WGS data.

Despite striking phenotypic diversity, both IDH-wildtype glioblastomas GB1 and GB2 displayed large truncal mutation burdens, characterized by clock-like mutational signatures SBS1 and SBS5, and unknown SBS40 (Supplementary Fig. 9). In GB1, most CNAs were clonal, including one-copy gains on chromosomes 7, 19, 20, and 21, focal high-level *EGFR* amplification (33–55 copies), and LOH on chr7q, 8p, 10, 15q, 16q, 18p, 19q, and 22, leading to hemizygous deletion of e.g. *PTEN*, while the remaining copies of chr7q and chr19q segments were duplicated in *MET* and *CCNE1* oncogene loci, respectively (Fig. 2b, Supplementary Fig. 5). The tumor suppressor *CDKN2A* locus on 9p was disrupted by rearrangements resulting in a nearly clonal homozygous deletion across all regions (Supplementary Fig. 7, Supplementary Tables 6 and 9). GB1 showed higher SV burden than the other tumors, most involved with the *CDKN2A* deletion and related rearrangements with chr11p, 18p, and 19q. Consistent with CNA architecture indicating that most genetic variation arose early, 73% of SNVs (4825 out of total 6621) were clonal (Supplementary Fig. 8), including full inactivation of *PTEN* (stop-gain p.R130X) (Supplementary Table 3). Two prominent late-arising subclones containing over 100 SNVs emerged at the opposite tumor poles, one specific to regions R3–R4 (purple) and containing *EGFR* frameshift (p.I1018Pfs*2 with 0.23 and 0.21 VAF indicating presence in 10 and 7 copies in R3 and R4, respectively), the other in regions R1–R2 (orange) with shared subclonal losses on chr1p, 9q, and 11p. These data indicate predominantly early and linear genomic evolution with late branching separating tumor poles in addition to pronounced heterogeneity in histological appearance.

GB2 showed a more divergent, but similarly trunk-dominated evolutionary pattern, characterized by high ploidy (4.66–5.83), variable total copy number per chromosome, and widespread LOH across chromosomes 4, 10, 12, 15, and 17, affecting tumor suppressor genes (*TP53* and *NF1* on chr17p, *PTEN* on chr10q, *KMT2D* on chr12q)(Fig. 2c, Supplementary Fig. 6, Supplementary Table 6). These LOH events occurred early during tumor evolution and were followed by multiple duplications across the whole genome, consistent with at least one WGD. The *EGFR* locus (7p) was massively amplified (>50 copies) in all regions, partly attributable to isochromosomes with two 7p arms, as observed in karyotyping (Supplementary Table 5). SNV clustering again revealed predominantly linear evolution, with 5852 (82%, total 7101 SNVs) truncal SNVs including mutations in *TP53* (p.R248Q), *PTEN* (p.I28T), *EGFR* (p.E317K), *RB1* (p.R418Efs*1) (Fig. 2c, Supplementary Fig. 8, Supplementary Table 4). Late-emerging subclones partitioned the tumor into two evolutionary branches, with nonsynonymous mutations in the left and right branch (e.g. *RELN* (p.C674S) gene, frequently altered in glioblastoma^23^). Tissue region GB2-R2 and cell line L3 were dominated exclusively by one branch, whereas other regions carried mixtures of both branches. Cell lines L1 and L2 were similar to R1 in mutational composition, which also contained the latest subclones among tumor regions. One subclone was detected only in L1 and L2, plausibly reflecting expansion of low-frequency variants during in vitro culture. Copy number profiles between cell line and tissue WGS samples were consistent, considering the technical discrepancies caused by absence of normal cell type admixture in cell lines (Supplementary Fig. 6). In addition, extensive LOH, high ploidy, and low NRPCC (<10), particularly in region GB2-R2 with estimated tumor purity of 23%, limit sensitivity for subclonal inference, warranting cautious interpretation of fine-scale branching. Overall, *TP53* mutation and these evolutionary characteristics suggest a possibility for an unstable tumor phenotype with transient genomic instability followed by ongoing copy number evolution^24,25^. Interestingly, many of the detected alterations (widespread copy neutral LOH, *TP53* and *RB1* mutations) are typical for giant-cell glioblastoma^26,27^, which is compatible with observed giant cells in GB2-R2 (Fig. 1c, Supplementary Fig. 2).

To further characterize the GB2 evolution in cell lines L1–L3, we utilized scRNA-seq to infer CNAs and subclones (Fig. 2d, Supplementary Fig. 11a,b). The single-cell data recapitulated the two main evolutionary branches identified by WGS and resolved nine additional subclones with higher granularity (Fig. 2e). While L1 yielded a substantially lower number of cells (Supplementary Table 1, Supplementary Fig. 12), majority of those mapped to right branch clones (3–5) together with L2, whereas most L3 cells mapped to left branch clones (7–11), consistent with the relationship between tissue regions R1 (right branch) and R2 (left branch). Chr16 loss, first detected in clone 3, corresponded to reduced total copy number in WGS of L1 and L2 and a subclonal deletion in region R1 (Supplementary Fig. 6). Descendant clones 4 and 5 consist almost exclusively of L2 cells with e.g. chr6 loss, consistent with the lower copy number observed in L2 WGS (and in L1, but underrepresented in single-cell data), whereas all other clones contain a mixture of cells from at least two lines. On the left branch, clone 7 was defined by chr1p amplification in scRNA-seq, but this amplification appeared truncal by WGS. In addition, a small chr11q deletion in clone 9 appeared clonal in L3 and subclonal in L2 WGS, mirroring the proportions seen in single-cell data. WGS also identified a subclone shared exclusively by L1 and L2 that could not be unambiguously resolved in the scRNA-seq data, likely due to the lower number of L1 cells. Despite a few platform-specific discrepancies—expected given differences in purity, ploidy, and resolution—scRNA-seq and WGS converged on a consistent underlying evolutionary topology.

### Genetic subclones occupy divergent transcriptional states shaped by regional microenvironment

To determine whether intratumoral heterogeneity is more pronounced at the transcriptional than at the genomic level and to assess how distinct genetic subclones partition into different transcriptional states, we analyzed bulk-RNA-seq from the same samples.

Across all tumors and regions, cancer cells predominantly exhibited an astrocyte-like (AC-like) transcriptional state (Fig. 3a)^15^. In addition, IDH-mutant DA1 displayed a higher proportion of neural progenitor-like (NPC-like) cells than the glioblastomas, whereas IDH-wildtype glioblastomas showed greater representation of hypoxia-independent mesenchymal-like (MES1-like) transcriptional states^21^. MES1-like states were particularly enriched in the peritumoral regions GB1-R1, and GB2-R1, although mesenchymal states are more typically found in tumor bulk or core areas^28,29^. Oligodendrocyte progenitor-like (OPC) activity (>10%) was observed only in GB1-R4 and GB2 regions R2 and R3.

**Fig. 3.**
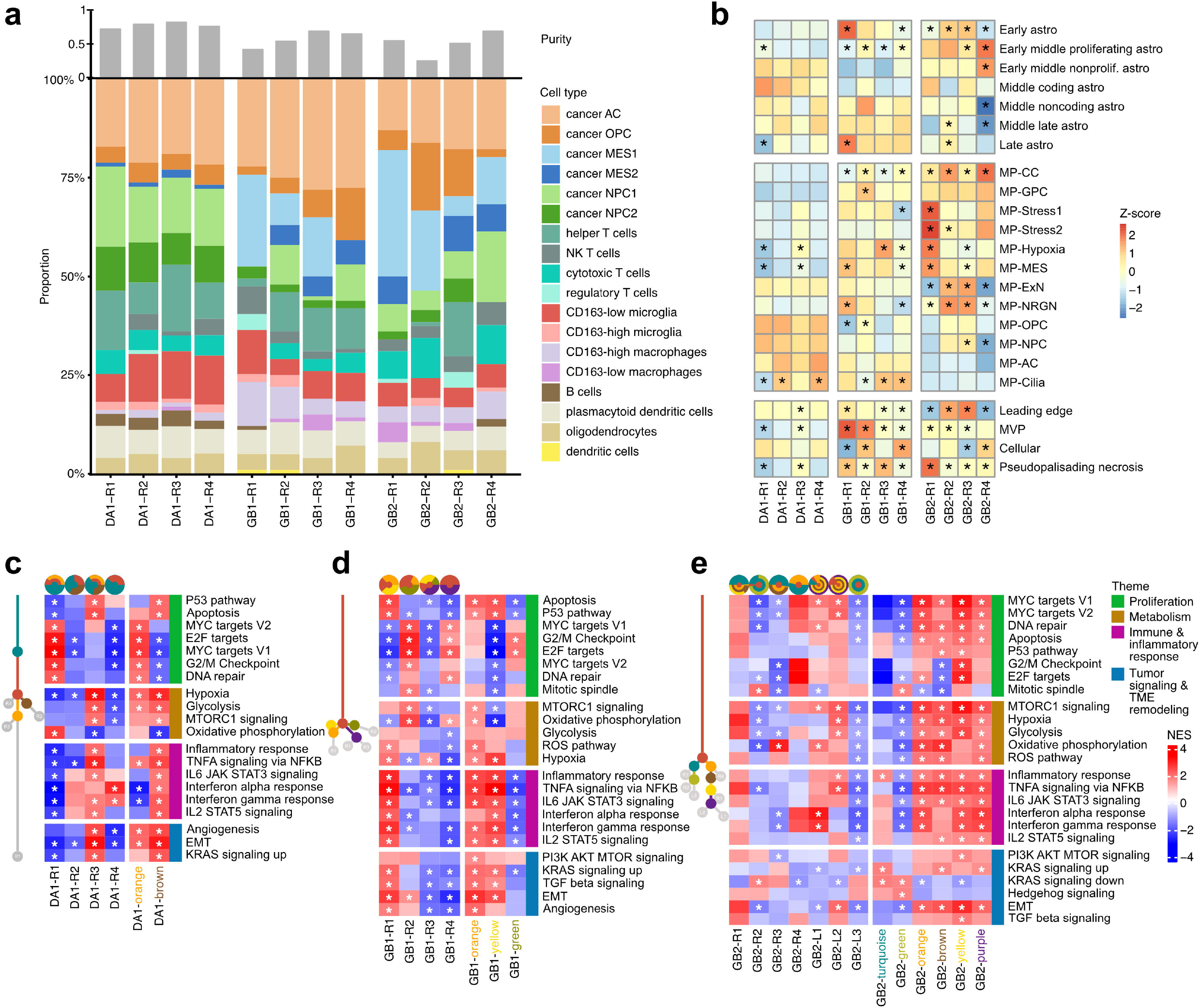
Bulk transcriptomic profiling reveals regional transcriptional divergence despite shared genetic subclones. **a)** Deconvolution of diffuse astrocytoma cell types from bulk RNAseq shows differences between IDH-mutant DA1 and glioblastoma tumors with regional heterogeneity. **b)** Activity z-scores of astrocyte maturation, glioblastoma metaprograms, and Ivy GAP gene sets across tumor tissue regions. Significant differences (Student’s t-test, p-value<0.01) of gene level z-scores between each region against other patient’s regions are shown with asterisks. **c-e)** GSEA of hallmark sets between regions (left) and subclone-enriched regions (right) in **c)** DA1, **d)** GB1, **e)** GB2. Jawbreaker plots show the subclonal compositions of each tumor region. Regions containing subclones were compared against the rest of the regions on the right. Significant hallmark sets relevant in proliferation, metabolism, immune and inflammatory response, and tumor signaling and TME remodeling are shown and marked with asterisks.

Mesenchymal programs were absent in the IDH-mutant DA1, and accordingly, macrophages were scarce^30^. Instead, DA1-R2–R4 were enriched for CD163-low microglia. In hypermutated DA1-R1, helper and cytotoxic T cells were elevated, whereas CD163-low microglia were reduced, indicating an increased T cell recruitment (Fig. 3a). Furthermore, gene set scoring and enrichment analysis based on differential expression of DA1-R1 demonstrated activation of proliferation, DNA repair, and oxidative phosphorylation, alongside the suppression of interferon signaling, leukocyte activation, cytokine signaling, and late astrocyte maturation, consistent with a rapidly dividing, metabolically active, and immunologically quiescent NPC-like component (Fig. 3b–c, Supplementary Fig. 13)^15,21,31^. In contrast, DA1-R3—despite sharing a WGS-defined subclone with R1 (Fig. 2a)—showed opposite activation of multiple signatures, including P53 and apoptosis, in addition to hypoxic, glycolytic, and inflammatory pathways typically associated with MES-like states^15,32^, but without MES-like cancer cells. This illustrates that regions sharing a genetic subclone can harbor cancer cell populations in distinct transcriptional states depending on the local microenvironment. To study the differences between samples in evolutionary context defined with WGS, regions were linked to genomic subclones based on enrichment of a specific clone in the sample. When comparing regions DA1-R1 and R3, containing a late-emerging orange subclone, against R2 and R4, representing earlier evolutionary phases, we saw enrichment for proliferation-associated E2F and MYC targets and DNA repair.

The GB1 peritumoral region R1, located at histologically apparent interface between less and more aggressive tumor areas, exhibited increased CD163-low microglia, CD163-high macrophages, and NK and regulatory T cells when compared to other regions, coexisting with abundant MES1-like cancer cells (Fig. 3a)^33,34^. This reactive edge state was characterized by immune activation, high inflammatory and interferon signaling, P53 pathway, and hypoxia combined with mesenchymal, proliferation-suppressed tumor cells (Fig. 3b,d, Supplementary Fig. 13), suggesting an intermediate mesenchymal state rather than strictly hypoxia or astrocyte-associated^34^. Both early and late astrocyte maturation programs showed high activity in R1, in line with the apparent tumor border with more and less differentiated histology (Fig. 1b). R1 histology exhibited gemistocytic differentiation, corresponding to immune and mesenchymal activation reported in IDH-mutant astrocytoma^35^. GB1-R3 displayed spindle cell differentiation, associated with mesenchymal stem cell-like glioblastoma phenotypes^36,37^, represented a different MES1-like subtype, but suppression of inflammatory, oncogenic, proliferative and metabolic pathways. GB1-R2 and R4 showed decreased MES1 and increased NPC1 and proliferation activities. GB1 subclone-associated regions displayed distinct transcriptional programs that mirrored the most dominant evolutionary branches (orange vs. purple, Fig. 2b). Branch-associated expression differences reflected microenvironment-responsive plasticity potentially driven by increased CD163-high macrophage involvement, epithelial–mesenchymal transition (EMT), and angiogenesis in the orange branch shared by GB1-R1 and R2 regions (Fig. 3d). Notably, regions R1 and R2 exhibited an increase in MVP signature^38^, which was also seen in their histology (Fig. 1c, Supplementary Fig. 2). The regionally distinct states, arising despite shared early truncal variants, illustrate that in GB1, genomic similarity coexists with substantial transcriptional plasticity across subclones.

GB2 peritumoral region R1 was dominated by MES1-like states combined with hypoxic signaling, along with metabolic, and pseudopalisading necrosis programs, representing a hypoxic mesenchymal niche around pseudopalisading necrosis seen in the tissue morphology^29,39,40^ (Fig. 3a–b,d, Supplementary Fig. 13). GB2-R4 showed the highest overall proliferation and early-to-mid astrocyte maturation signatures, together with elevated NPC1-like fractions. Tumor core regions GB2-R2 and R3 showed neuronal gene set activities and low proliferation and immune signals, in addition to relatively low tumor purity especially in R2; extensive necrosis as detected based on histology likely contributes to both reduced proliferation and lower purity estimates. Transcriptomic differences between regions enriched for distinct genomic subclones aligned with the evolutionary branches identified by WGS (Fig. 2c). The right-side branch subclones, many of which appeared in cell lines, showed increased inflammatory, hypoxic, EMT, and mitotic activity. In contrast, the left branch was dominated by Hedgehog signaling. Overall, GB2 exhibited the strongest proliferation signature, consistent with its high Ki-67 index (∼40%) and densely cellular morphology (Fig. 1c, Supplementary Fig. 2).

### Single-cell profiling reveals subclone-associated and niche-dependent cell state heterogeneity in GB2

To validate and extend the transcriptional heterogeneity observed in bulk RNA-seq, we profiled the transcriptomes of three GB2 neurosphere lines L1–L3.

Single-cell transcriptomes revealed clear region-linked differences in cancer cell states. L1 and L2 consisted almost exclusively of MES1-like cells (Fig. 4a), whereas L3 retained a more heterogeneous composition spanning MES1-, NPC1-, and AC-like states. These patterns closely mirrored the tissue bulk deconvolution profiles and aligned with subclonal relationships: GB2-R1, enriched for MES1 states in bulk RNA-seq, corresponded to the MES1-dominated L1 and L2 cultures. Consistently, L1, L2, and R1 shared an evolutionary branch marked by alterations such as chr16 loss detectable in both scRNA-seq and WGS data, and scRNA-seq-derived clones enriched for L2 cells (>50%; clones 4, 5, and 12) were dominated by MES1-like programs (Fig. 4b–c, Supplementary Fig. 11c). In contrast, GB2-R2, which displayed a mixture of AC, OPC, MES1, and NPC states, retained both state distribution and subclonal structure in the L3 neurosphere (Fig. 3a, Fig. 4a). The corresponding scRNA-seq clones populated primarily by L3 cells (clones 7, 8, 9, and 11) belonged to a distinct evolutionary branch characterized by chromosome 1 gains and exhibited a more diverse mixture of NPC-, AC-, and MES-like states (Fig. 4b–c, Supplementary Fig. 11c). This indicates that the neurosphere cultures largely preserve region-specific transcriptional states and subclonal structure of their tissue of origin, despite adaptation to culture conditions and continued genomic evolution occurring during in vitro expansion.

**Fig. 4.**
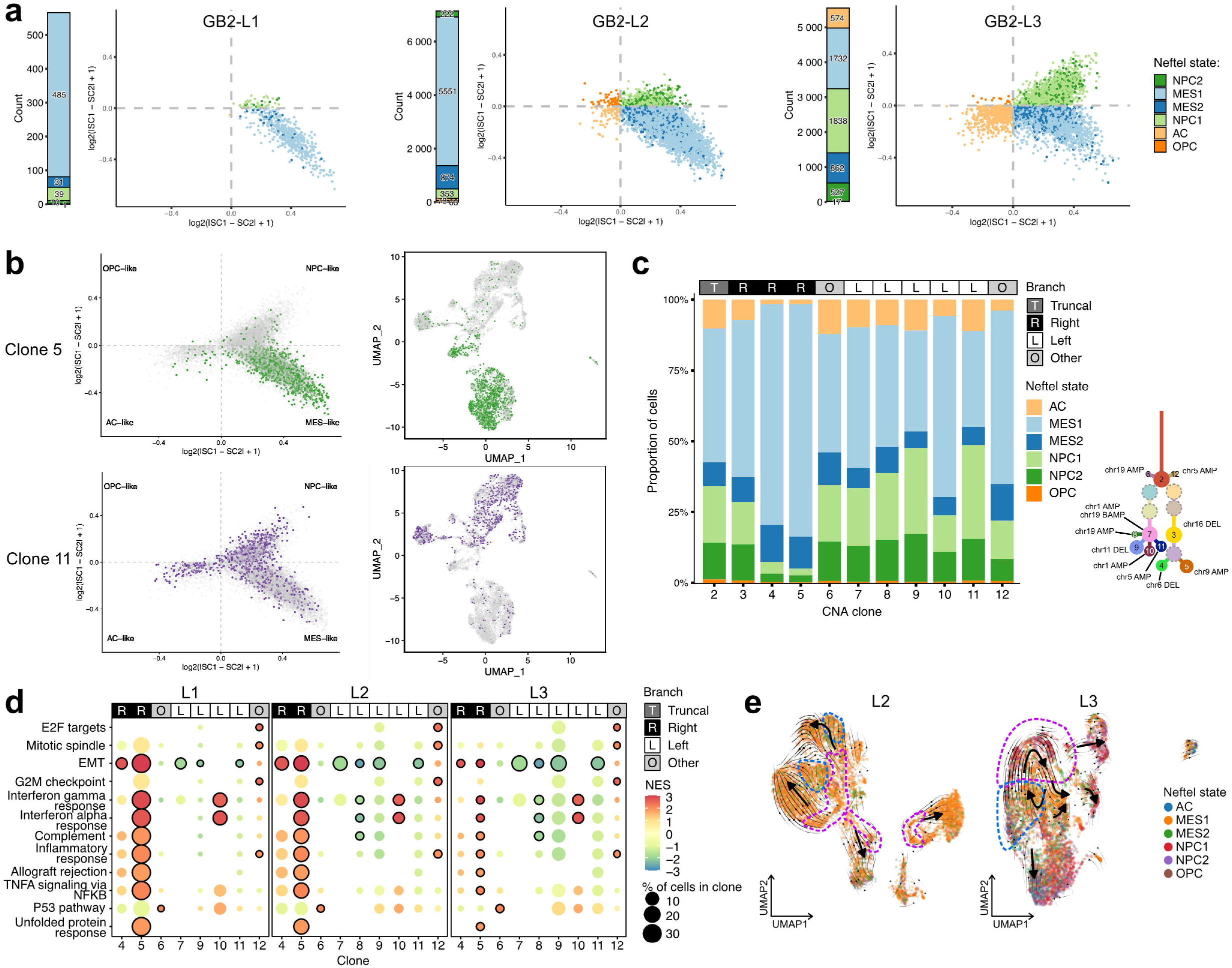
Single-cell profiling of GB2 cell lines reveals subclone-associated transcriptional states and dynamic cell-state transitions. **a)** Glioblastoma cell states in scRNA-seq of GB2 cell lines. L1 and L2 represent mostly MES1-like cells, whereas L3 cells are also in NPC and AC states. **b)** Clone 5 (descendant of the yellow WGS clone on the right branch) and clone 11 (descendant of the green WGS clone on the left branch) represent different branches and cell states in scRNA-seq CNA-based tumor evolution in integrated neurosphere cells. **c)** CNA clone glioblastoma state compositions follow cell line sample proportions and CNA evolution branching. **d)** Hallmarks GSEA between scRNA-seq CNA derived clones showed globally elevated transcriptional activity (clones 5, 4, 10, and 12) or quiescent profiles (clones 7, 8, 9, and 11). Only the clones that have significant (circled, based on Wilcoxon rank-sum test, adjusted p-value<0.05) enrichments are shown. **e)** RNA velocity flow reveals low-proliferating endpoints in L2 and L3 and dynamic transitioning between cell cycle phases in L3. Cells in S-phase and G2/M-phase are marked with blue and purple dashed lines, respectively.

Differential expression across scRNA-seq-defined clones demonstrated marked functional heterogeneity independent of cell-cycle activity. Clones 5 and 4 representing the right evolutionary branch and clones 10 from left and 12 from a separate short branch showed globally elevated transcriptional activity, whereas clones 7, 8, 9, and 11 representing the left branch displayed reduced expression and quiescent profiles (Fig. 4d). Transcriptionally highly active clones were enriched for either proliferation (clone 12) or immune-hot, mesenchymal, inflammatory, and stress-adaptive signatures with reduced cycling, consistent with the predominance of MES1-like states in these clones^15,32^. When restricting analysis to cell-line-intrinsic heterogeneity, proliferative populations formed distinct clusters in L2 and L3, while less-cycling clusters were enriched for inflammation pathways (Supplementary Fig. 14–15).

RNA velocity analysis highlighted dynamic trajectories. In L2’s proliferative clusters, particularly an inflammation-associated (interferon, TNFA via NFKB, and hypoxia) cluster 6 and G2/M-phase cluster 8, were positioned upstream of several low-cycling states (clusters 1 and 4), including a hypoxia-retaining state (cluster 0) (Fig. 4e, Supplementary Fig. 15–16). Cluster 6 transitioned toward cluster 0 via an intermediate proliferating population. A separate trajectory (cluster 10) was enriched for angiogenesis signature. In L3, dynamic cycling transitions occurred between low-proliferative inflammatory MES-like states (clusters 4 and 8), proliferative S-phase (clusters 2 and 5), and G2M NPC-like state (cluster 3). These proliferative and reactive populations appeared to diverge toward low-proliferating endpoints, including a quiescent AC-like population (cluster 0) and NPC-like states (clusters 6 and 7).

Across tumors, transcriptomic heterogeneity was substantially more pronounced and spatially structured than genomic heterogeneity, with region-specific differences in cancer cell-state composition, stress and inflammatory programs, and proliferation signatures emerging within largely shared genetic backbones. In GB2, transcriptomic similarities between spatial regions and their matched cell lines frequently aligned with genomic relationships established by subclonal reconstruction, indicating cross-modal consistency without implying genetic determinism of cell-state identity. Together, these observations highlight the coexistence of conserved genomic architecture with spatially variable, microenvironment-associated transcriptional states.

## Discussion

This multi-region whole-genome and transcriptomic analysis provides an integrated view of how genetic evolution and transcriptional plasticity jointly shape intratumoral heterogeneity in high-grade diffuse astrocytomas. Across two glioblastomas and one IDH-mutant grade 4 astrocytoma, WGS revealed predominantly early and linear evolutionary trajectories, characterized by extensive truncal backbones and late subclonal branching in glioblastoma and a comparatively shorter trunk with additional regional diversification in the IDH-mutant astrocytoma, including a localized hypermutation in one region. In contrast, transcriptomic profiling uncovered pronounced spatial variation in cancer cell states and pathway activity that aligned with microenvironmental context and region–cell-line pairing, often consistent with, but not dictated by, genomic relationships. Despite the overall limited genomic divergence beyond truncal events, we observed individual or regionally restricted genetic alterations that were partly associated with phenotypic differences. These findings support a model in which clinically actionable genomic alterations are largely truncal, whereas phenotypic heterogeneity is primarily plastic and driven by niche-dependent programs rather than ongoing genetic diversification.

This distinction has important clinical implications. While single biopsies may capture truncal, actionable genomic drivers, the spatially variable distribution of phenotypic states and therapeutic targets, such as EGFR amplification, likely contributes to treatment failure^41^. Resolving the full spectrum of functionally relevant heterogeneity will therefore require integration of genomic and state-level profiling across multi-region and single-cell approaches.

Our findings refine previous models of glioma evolution. Most diffuse glioma tumor evolution studies focus on recurrence^21,42–44^, and primary tumor spatial heterogeneity, particularly in IDH-mutant astrocytomas, remains less characterized. Multi-regional exome sequencing studies have reported regional variation in driver alterations in gliomas^8,11,12,14^, but limited genomic coverage may overestimate the extent of heterogeneity^45^. In contrast, our WGS data indicate largely truncal driver alterations across all tumor regions, suggesting that much of the genomic architecture underlying tumor progression is established early^46^. To our knowledge, multi-regional WGS-based evolutionary analysis has been reported for only one diffuse glioma (retrospectively in an IDH-mutant astrocytoma)^47^.

We detected both spatial branching towards opposite tumor poles (GB1), and intermixing of later-phase subclones (DA1 and GB2)^8^. In addition, the spatial heterogeneity of each tumor was driven by different large-scale genomic architectures; transient genome-wide genomic instability and high ploidy resulting from punctuated evolution^24,25^ (GB2), *CDKN2A* inactivation-associated structural remodeling^43,48^ (GB1), or chromothripsis, rarely occurring in IDH-mutant tumors, in addition to WGD (DA1)^8,49^. GB2’s histologically aggressive features corroborated its genomic complexity.

Regardless of the relatively constrained genomic divergence, tumors harbored substantial spatial variation in transcriptional subtypes^1,11,14^. Glioblastoma cell states were strongly influenced by microenvironment conditions and diverged from hypoxia-independent MES1-like at peritumoral sites to a mixture of NPC1- and OPC-like states in deeper regions despite largely shared genetic backgrounds. Although all peritumoral glioblastoma regions appeared as tumor-normal interfaces intraoperatively and radiographically and harbored some of the latest evolutionary subclones^50^, they exhibited high tumor cell content, mesenchymal activity, histological core features and proliferation comparable to tumor core, unlike the typical peritumoral zone^28,29,38,51^. This suggests substantial infiltration beyond radiographic margins, highlighting their limited ability to reliably delineate tumor extent^52–54^. These findings emphasize the importance of considering microenvironment-driven plasticity when interpreting spatial tumor organization.

Spatially organized histologic phenotypic differences arose within tumors sharing a common genomic driver landscape. Although histological phenotypes were consistent with Ivy Glioblastoma Atlas Project (Ivy GAP) signatures^38^, they did not map to genomic subclones but instead corresponded variably to transcriptional states: pseudopalisading necrosis to hypoxia and MES2 state, and MVP to proliferation and immune cell domination, and leading edge to neuronal metaprograms, albeit in GB2 tumor core rather than peritumoral regions^21,29^. Together, these findings suggest that morphological features, including pseudopalisading necrosis and MVP, and differentiation patterns likely arise from environment-driven cellular states rather than underlying genetic differences.

The observation of localized hypermutation in DA1-R1 further illustrates the temporal decoupling between genomic evolution and phenotypic manifestation, as the region appeared more advanced genomically than its grade 3 histology. Notably, the accumulation of mutations may also promote a more diffuse, infiltrative growth pattern instead of inducing the morphological hallmarks required for grade 4 classification, such as necrosis or MVP. Genomic changes can accumulate without immediate phenotypic consequences, and conversely, pronounced phenotypic divergence can occur in the absence of substantial genetic differentiation, reinforcing intratumoral heterogeneity as interplay between relatively stable genomic scaffold and highly dynamic, environment-responsive transcriptional programs^2^.

Several factors merit consideration. The cohort size (n=3) enabled deep multi-modal profiling but limits generalizability. Tumor purity and ploidy influence the sensitivity of subclonal inference, particularly for GB2, and scRNAseq may diverge from WGS in highly polyploid contexts^55^. Patient-derived cultures can further shift state compositions and expand rare subclones during in vitro propagation. While single-cell analysis of the neurosphere cultures recapitulated the major WGS-defined evolutionary branches, it also revealed additional minor subclones, suggesting ongoing copy-number selection during the 13–16 in vitro passages. Despite this, cell lines largely preserved both the genomic and transcriptomic features of their matched tissue regions, including cell-state compositions and transitions along the proneural-mesenchymal axis as inferred by RNA velocity^56^. Across clones, cells segregated into proliferative and inflammatory modules, supporting a model in which glioma cell states exist along dynamic continuums shaped by both lineage and microenvironment.

Across all tumors, truncal driver mutations were shared, but distinct genetic subclones were associated with different transcriptional states across spatial regions. Conversely, genetically similar regions can diverge transcriptionally under local microenvironmental pressures. Thus, intratumoral phenotypic diversity arises from a combination of subclone specific programs and context dependent plasticity rather than ongoing acquisition of region specific driver events. Transcriptomic similarities between spatial regions and their matched cell lines often paralleled subclonal relationships, reinforcing internal consistency without implying genetic determinism of cell state identity. Histological features such as MVP, pseudopalisading necrosis, or gemistocytic differentiation similarly represented convergent morphologic outcomes emerging from distinct underlying states. Overall, these results highlight that spatial heterogeneity in high grade astrocytomas is driven largely by transcriptional diversification superimposed on shared genomic scaffolds, underscoring the need for multimodal regional sampling, deep whole genome sequencing, and high resolution single cell or spatial transcriptomic approaches to fully resolve the functional diversity that shapes clinical behavior.

## Methods

### Sample cohort

The study design was approved by the Regional Ethics Committee of Pirkanmaa Hospital District (now the Ethics Committee of the Wellbeing Services County in Pirkanmaa; approval number R14024). Tumor samples had been obtained from surgery patients at Tampere University Hospital and diagnosed according to the World Health Organization (WHO) 2021^57^ classification for this study. Three large and aggressive tumors were included in the study, one IDH-mutant grade IV diffuse astrocytoma (DA1) and two IDH-wt glioblastomas (GB1 and GB2) (Supplementary Fig.1). Samples from four spatially distinct tumor loci were collected from each tumor: one locus from the peritumoral region (R1), two from the centroid of the tumor (R2 and R3), and the fourth one from the opposing tumor side (R4) (Supplementary Fig.2). Fresh frozen samples from each loci were embedded in OCT cryomount (Tissuetek, Sakura Finetech, Japan) and frozen at -80. Blood samples were collected pre- or postoperatively in EDTA tubes and preserved at -80. In addition, patient-derived neural stem cell cultures were established as described below from freshly resected tumor tissue of one patient, obtained from two regions adjacent to the frozen samples of the same patient.

### Establishment of patient-derived cell cultures

Two different loci from GB2 were processed into cell lines directly from the operating room. The first loci was processed into two cell lines (L1 and L2) and the second loci into one cell line (L3). Necrotic and burnt tissue along with large blood vessels were manually removed. The tissue was minced with a scalpel and tumors were enzymatically digested using Tumor dissociation kit (#130-095-929, Miltenyi Biotech). Acquired single-cell suspension in DMEM/F12 (#21331, Gibco) was filtered through a 70 µm cell strainer, pelleted and resuspended in neural stem cell media (DMEM/F12, DMEM Neurobasal medium (#21103049, Gibco), B-27 supplement (#17504044, Gibco) and N-2 supplement (#17502048, Gibco)). Cells were cultured for several days to allow neurospheres to form. After sphere formation, the spheres were transferred to laminin-coated (#L2020, Sigma-Aldrich) Primaria plates (Corning) and maintained after that as monolayer cell cultures for 14 (L1), 16 (L2), or 13 (L3) passages.

### RNA and DNA extraction and sequencing

Fresh frozen tissues were sectioned into 10 µm slices using a cryomicrotome with alternating sections used for RNA and DNA extraction, respectively. Cell lines were harvested with scraping over ice and pelleting after that. Genomic DNA was extracted using QIAmp Mini kit (#51304, Qiagen, Hilden, Germany). Messenger RNA was extracted with mirVana miRNA isolation kit (#AM1560, ThermoFisher Scientific, Waltham, MA, USA). Germline DNA was extracted as a control form EDTA blood samples using QIAamp DNA Blood Mini Kit (#51304, Qiagen). Library construction and sequencing of RNA and DNA were performed by Novogene (Beijing, China). DNA sequencing libraries were generated using NEBNext Library Prep Kit (E7103S, New England Biolabs), PCR enriched by P5 oligos, indexed by P7 oligos and purified with AMPure XP (A63881, Beckman Coulter). For RNA sequencing library preparation, mRNA was first fragmented with a fragmentation buffer and then synthesized into cDNA with hexamer primers, synthesis buffer (Illumina), dNTPs, RNase H and DNA polymerase I. Then, after a series of terminal repair, ligation and adaptor ligation, the double stranded cDNA library was completed through a size selection and PCR enrichment. Sequencing for both RNA and DNA was done with Illumina NovaSeq 6000 with paired-end 150 bp strategy (PE150). The targeted sequencing depth for whole-genome sequencing was 30x for blood-derived and cell line samples and 60x for tumor tissue samples (Supplementary Table 1).

### Single-cell RNA sequencing

Patient-derived cell lines were subjected to single-cell RNA sequencing with the Chromium library preparation method (10x Genomics, Pleasanton, CA, USA). Cells were harvested by detachment with Accutase (A1110501, Gibco), pelleted (1100 rpm, 5 min) and washed once with PBS. Cells were counted with LUNA automated cell counter (Logos Biosystems, South Korea) and 20,000 cells were loaded to Chromium microfluidics chip (#1000127, 10x Genomics). Library preparation was carried out with Chromium™ Single Cell 3’ Library & Gel Bead Kit v2 (#1000269, 10x Genomics) according to the manufacturer’s recommendations. Sequencing was performed at Novogene (Beijing, China) with Illumina NovaSeq 6000 technology paired-end 150 bp strategy (PE150) (Supplementary Table 1).

### Karyotyping

Cell lines L1–L3 were plated on a laminin coated primaria plates and cultured in neural stem cell media for 7 days. During the culturing, cell lines were passaged twice. Cells were then subjected to karyotyping, which was performed as a service in Fimlab (Tampere University Hospital, TAYS) using routine G-banding at 300-400- band per haploid chromosome set resolution following the International System for Human Cytogenomic Nomenclature (ISCN) 2020^58^ (Supplementary Table 5).

### WGS analysis

Whole-genome sequence reads were aligned to human reference genome GRCh38 using BWA-MEM v0.7.17. Duplicate reads were marked with samblaster v0.1.24, BAM files sorted with samtools v1.8, and read-groups were assigned using Picard v2.21.8. Downstream processing followed GATK Best Practices for somatic short variant discovery, including Base Quality Score Recalibration (BQSR) performed with GATK v4.1.1.0 using known sites from the GATK Resource Bundle (1000G_phase1_snps, Mills_and_1000G_gold_standard_indels, Homo_sapiens_assembly38_dbsnp138). Somatic single nucleotide variants (SNV) and insertions and deletions (indels) were detected from tumor and cell-line WGS data with GATK Mutect2 using genomic DNA isolated from leukocytes (blood) from each patient as a matched normal and GATK Best Practices Bundle AF-only gnomAD as germline resource file containing population allele frequencies of common and rare variants. Mutect2 was run with a panel of normals (PoN) consisting of 62 blood WGS samples from the International Cancer Genome Consortium (ICGC)^59^ that were aligned to GRCh38 and processed through the variant calling workflow in a similar manner to our patient WGS data, and their tumor-only variant calls were combined using GATK v4.0.11.0 CreateSomaticPanelOfNormals. The detected variants were filtered using GATK FilterMutectCalls and bcftools v1.8. Annovar (v2019Oct24) was used for the annotation of the SNVs (Supplementary Table 2–4) using the following hg38 databases provided by the Annovar website: RefSeqGene v20190929, ExAC 65000 exome allele frequency data v0.3, dbNSFP v4.1c, gnomAD whole-genome data v3.0, and the COSMIC GRCh38 v92 database not included in the Annovar package. Germline variant calling was performed with GATK version 4.2.5.0 HaplotypeCaller, CNNScoreVariants, and FilterVariantTranches using default tranche values for SNPs (99.95) and indels (99.4), and annotated with Funcotator v1.6.20190124g and/or Annovar RefSeqGene 20190929, avSNP v150, dbNSFP v4.2c, gnomAD v3.12, and ClinVar v20221231. Cosmic signature presence was analyzed with SigProfilerAssignmentR v.1.1.3 wrapper^60^.

Structural variants (SVs) were identified with SvABA v1.1.3^61^ from tumor and cell line sample WGS BAM files using the corresponding patient blood DNA as normal and GATK Resource Bundle known indels as a reference database. SVs were annotated with a script provided in SvABA github (Supplementary Tables 8–10). Copy number alterations (CNAs) were analyzed using Battenberg v2.2.10^62^ with provided GRCh38 reference files for matched tumor and blood normal sample WGS BAM files. alleleCounter v. 4.0.2, Beagle 5.0 v12Jul19, and copynumber v1.26.0 were used with Battenberg for allele counting, phasing, and segmentation, respectively, and breakpoints detected with SvABA were included in initial segmentation. In-house software cn_solver was used to get additional estimates of purity and ploidy (Supplementary Table 1), which were used in refitting final Battenberg solutions (Supplementary Table 6–7). In addition, for patient GB2, karyotype analysis from cell line samples was used for validating purity and ploidy estimation.

### Tumor evolution analysis

Subclonal reconstruction was performed using the multidimensional DPClust^62^ method (https://github.com/Wedge-lab/multidimDPClust) by clustering somatic SNVs detected in the tumor and cell line samples based on their sample-wise cancer cell fractions (CCF); VAF scaled by tumor purity and local copy number from Battenberg (Supplementary Table 2–4). Cluster median CCF values were estimated by fitting a left-truncated binomial distribution curve on the cluster SNV CCFs as described in^63^. Clusters subclonal in all samples with smaller median CCFs than descendant clusters and clusters shared by two samples containing fewer than 0.5% of total SNVs were excluded. Evolutionary trees were manually derived from non-conflicting median CCF values based on pigeonhole principle. Due to high ploidy and low NRPCC in patient GB2 samples, conflicting cluster median CCFs were resolved based on other samples and minimal evolution principle. Driver alterations associated with glioblastoma and diffuse astrocytoma (Supplementary Table 11) were assigned to the evolutionary clusters by their CCF. Private clusters were interpreted as descendants of clones with most similar median CCF in evolutionary trees. Jawbreaker visualizations^63^ were generated to show the evolutionary lineages of each clone and their proportions on the layered circle sectors and arcs.

### Bulk RNA-seq analysis

Read alignment was performed with STAR v2.7.10a against the hg38 reference genome (GENCODE release 40). Genewise read counts were determined with featureCounts v1.6.2. Median of ratios normalization was applied (Supplementary Table 12). R-package DESeq2 version 1.40.2 with lfcShrink was used to determine the differentially expressed genes between each spatial location compared to other spatial locations within the patients. Also comparisons of clone specific spatial locations against other locations within the patient were performed. R-package fgsea version 1.26.0 was used to calculate the enrichment scores for Hallmark, GO and Kegg gene sets.

Deconvolution was performed with CIBERSORTx estimating the abundance proportions of different cell types from bulk RNA-seq samples. References of cell types were constructed with single-cell RNA-seq data from^15,16,64,65^. Cells were annotated and reference set was built as described in^66^ from following cell types: cancer cells NPC1, cancer cells NPC2, cancer cells OPC, cancer cells AC, cancer cells MES1, cancer cells MES2, CD163-low microglial cells, CD163-high microglial cells, CD163-low macrophages, CD163-high macrophages, dendritic cells, helper T cells, NK T cells, T regulatory cells, B cells, plasmacytoid dendritic cells, oligodendrocytes, and cytotoxic T cells. Proliferating cancer and myeloid cells were excluded. Combined activity z-scores^67^ were calculated for literature gene sets^7,15,16,38,43,68,69^ and significance was tested with Student’s t-test comparing gene level z-scores between one tumor region against others.

### Single-cell RNA-seq analysis

Cells were aligned and UMIs counted with Cell Ranger v7.1.0 against human reference genome GRCh38-2020-A provided by 10X Genomics. GB2-L1 sample yielded lower total cell count than other cell lines (796 cells compared to 8,626 in L2 and 6,251 in L3), which resulted in higher number of reads per cell (mean 309,023 versus 28,876 and 32,892 in L2 and L3, respectively). Overall data quality in terms of background separation and mapping percentages was desirable (Supplementary Table 1, Supplementary Fig. 12). CellBender v0.2.1 did not show signs of ambient RNA. Sample L1 cells were filtered for maximum mitochondrial DNA percentage of 20%, minimum 5000 genes, and expected doublet rate of 0.66% (according to 10X kit multiplet rate) per cell utilizing R package SCP v0.5.1 CellQC tool with scDblFinder method for doublet calling. Corresponding numbers for L2 cells were maximum 12% mitochondrial DNA, minimum 1500 genes, and 6.6% doublet rate, and for L3 sample cells: maximum 8% mitochondrial DNA, minimum 1500 genes, and 4.8% doublet rate. Seurat v5.3.0 R library was used for normalization, feature selection, clustering with resolution 0.8, and visualizations. Cell states were estimated using Neftel ^15^ glioblastoma gene sets in the Seurat AddModuleScore function and selecting the maximum score per cell. Cluster enrichments in MSigDB Hallmark gene signatures were analysed with SCP R library functions RunDEtest and RunGSEA. Additional selected gene set scores were calculated using AddModuleScore. The RNA velocities of single cells were estimated with scVelo v0.3.3 using loom files produced with velocyto v0.17.17 and metadata extracted from individual Seurat objects.

Copy numbers in single cells were estimated using Numbat v1.5.1^70^, R v4.4.0 for which germline SNPs were phased using Eagle v2.4.1 and their allele counts were calculated with cellsnp-lite v1.2.3 and bcftools v1.15.1. Single-cell raw RNA read counts from each cell line were integrated with Seurat function IntegrateLayers using Harmony v.1.2.4 method for Numbat analysis, normal cell expression reference was generated from publicly available glioblastoma scRNA-seq datasets^15,16,64,65^ by integrating normal cells identified as previously described^66^ with Harmony using 2000 variable features and 50 PCs and extracting raw counts. Battenberg cell line sample segments were inputted as consensus CNAs.

### Statistical tests

Significance in gene set z-score activities was tested with Student’s t-test comparing gene level z-scores between one tumor region against others. Differential expression between scRNA-seq-derived clones and clusters was assessed using Wilcoxon rank-sum test. Significant hallmark enrichments in scRNA-seq-derived clones and clusters were also estimated using Wilcoxon rank-sum test. Adjusted p-values were calculated using Bonferroni correction for scRNA-seq differential expression analysis and Benjamini-Hochberg false discovery rate for scRNA-seq enrichment analysis. All statistical tests were two-sided.

## Data availability

The datasets generated and/or analysed during the current study will be available in the Finnish Federated European Genome-phenome Archive (FEGA) repository once this has been approved based on the legal assessment and all the needed agreements have been made. The access to the data can be requested from the Research Services (Tutkimuspalvelut) of the data controller, the Wellbeing Services County of Pirkanmaa. Bulk RNAseq counts can be downloaded from GEO with an accession number GSE334421.

## Code availability

The code written for the analysis of the data presented in this publication are available in https://github.com/NykterLab/glioma-heterogeneity/ and https://github.com/nurmians/vcf_helper.

## Acknowledgements

We highly appreciate Paula Kosonen, Hanna Selin, Päivi Martikainen, and Marja Pirinen for their valuable technical assistance and Sofia Keitaanniemi and Henna Kujanen for their excellent help in computational analysis during the project. Furthermore, we want to deeply acknowledge the patients involved in this study and the personnel at Tampere University Hospital for their assistance in patient recruitment and Fimlab laboratory services for conducting the immunohistochemical staining for the diagnosis of the tumors as well as everyone in these institutes who has contributed to the sample collection. The authors acknowledge the CSC - IT Centre for Science, Finland, for providing computational resources and the Biocenter Finland (BF) and Tampere Genomics Facility for the service.

## Author contributions

S.O.: computational and statistical analysis of sequencing data, data integration, result interpretation, manuscript figures, writing the manuscript. S.M.: sample handling, result interpretation, writing the manuscript. R.N.: computational and statistical analysis of RNA sequencing data, involved in manuscript writing. I.H.: patient cell line establishment. A.N.: computational analysis of whole-genome sequencing data. A.M.T.: histological staining analysis, writing the manuscript. I.S.: computational image analysis, result interpretation. E.V.: sample handling, data generation. K.N.: clinical expertise. H.H.: sample collection, histological staining analysis, neuropathology and other clinical expertise. K.J.R.: design of the study, histological staining analysis, result interpretation, supervision. J.H.: design of the study, sample and clinical data collection, clinical expertise. M.N.: design of the study, data management, result interpretation, supervision, project coordination. All authors have reviewed and accepted the manuscript.

## Competing interests

The authors declare no competing interests.

## Supplementary Information

Supplementary Figures 1–16 can be found in Supplementary Figures.pdf file.

Supplementary Tables 1 and 5–12 can be found in a file Supplementary Tables.xlsx and Supplementary Tables 2–4 in a file Supplementary Tables (2–4).xlsx.

## Ethics approval and consent to participate

This study received approval from the Regional Ethics Committee of Pirkanmaa Hospital District (now the Ethics Committee of the Wellbeing Services County in Pirkanmaa; approval number R14024). Written informed consent was obtained from all participants. All ethical procedures complied with University of Tampere regulations and Declaration of Helsinki.

## Consent for publication

All participants of this study provided informed consent for publication of their data according to legislation and good practice.

## Funding

The study was financially supported by the Academy of Finland (#352818 to M.N.), Cancer Foundation Finland (M.N., S.O.), Pirkanmaa Cancer Society (M.N.), Sigrid Jusélius Foundation (M.N., K.J.R.), Competitive State Research Financing of the Expert Responsibility area of Tampere University Hospital (M.N., K.J.R., J.H., H.H., K.N.), Finnish Cultural Foundation (S.O., E.V.), The Paulo Foundation (S.O.), Ida Montin Foundation (S.O.), The Finnish Foundation for Technology Promotion (S.O.), Danish Data Science Academy (S.O.), Orion Research Foundation (S.O.), Emil Aaltonen Foundation (S.M.), and Tampere University Faculty of Medicine and Health Technology (S.O., S.M.)

## Notes

### Competing Interest Statement

The authors have declared no competing interest.

